# Microbial Diversity in a Military Impacted Lagoon (Vieques, Puerto Rico) as Revealed by Metagenomics

**DOI:** 10.1101/389379

**Authors:** Lizbeth Dávila-Santiago, Natasha DeLeón-Rodriguez, Katia LaSanta-Pagán, Janet K. Hatt, Zohre Kurt, Arturo Massol-Deyá, Konstantinos T. Konstantinidis

## Abstract

The Anones Lagoon, located in the island municipality of Vieques, Puerto Rico (PR), received extensive bombing during military practices by the US Navy for decades. After military activities ceased in 2003, the bombing range was designated as part of a larger Superfund site by US EPA. Here, we employed shotgun metagenomic sequencing to investigate how microbial communities responded to pollution by heavy metals and explosives at this lagoon. Sediment samples (0-5 cm) from Anones were collected in 2005 and 2014 and compared to samples from two reference lagoons, i.e., Guaniquilla, Cabo Rojo (a natural reserve) and Condado, San Juan (PR’s capital city). Consistent with selection under low anthropogenic impacts, Guaniquilla exhibited the highest degree of diversity with lower frequency of genes related to xenobiotics metabolism among the three lagoons. Notably, a clear shift was observed in Anones, with *Euryarchaeota* becoming enriched (9% of total) and a concomitant increase in community diversity, by about one order of magnitude, after almost 10 years without bombing activities. In contrast, genes associated with explosives biodegradation and heavy metal transformation significantly decreased in abundance in Anones 2014 (by 91.5%). Five unique population genomes were recovered from the Anones 2005 sample that encoded genetic determinants implicated in biodegradation of contaminants. Collectively, these results provided new insights into the natural attenuation of explosive contaminants by the benthic microbial communities of the Anones lagoon and could serve as reference points to enhance bioremediation actions at this site and for assessing other similarly impacted sites.

**Importance:** This study represents the first assessment of the benthic microbial community in the Anones Lagoon in Vieques, Puerto Rico after the impact of intense pollution by bombs and unconventional weapons during military training exercises. Evaluating the microbial diversity of Anones, represents an opportunity to assess the microbial succession patterns during the active process of natural attenuation of pollutants. The culture-independent techniques employed to study these environmental samples allowed the recovery of almost complete genomes of several abundant species that were likely involved in the biodegradation of pollutants and thus, represented species responding to the strong selection pressure posed by military activities. Further, our results showed that natural attenuation has proceeded to a great extend ten years after the cease of military activities.

## Introduction

Military exercises have left a legacy of pollution worldwide and represent one of the most substantial anthropogenic disturbances of natural ecosystems. Activities such as naval maneuvers and arms testing have resulted in the contamination of air, water, and soil with heavy metals and explosive compounds (1, 2, 3, 4, 5). In cases where military bases or ranges have been established in natural areas, a wide range of environmental damages has been documented (2, 3, 6, 7). At the moment, little is known about the magnitude of contamination by military training activities since a full-disclosure of what type and quantity of contaminants were used in any given location is typically unavailable. This leads to a potentially serious environmental hazard since explosive compounds cause not only physical damage of the environment where they detonate but also may have long-term consequences due to trace contamination or bi-products that can be arrested in soil, sediments, water, and even leak in groundwater (3, 8). Two of the most commonly used nitramine explosives in military activities since World War II is hexahydro-1,3,5-trinitro-1,3,5-triazine (RDX) and 2,4,6-trinitrotoluene (TNT). Both are categorized as carcinogenic by the Environmental Protection Agency (US EPA). Other explosives commonly used are octahydro-1,3,5,7-tetranitro-1,3,5,7-tetrazocine (HMX) and 2,4-dinitrotoluene (2,4-DNT) (3, 8). In general, insufficient information is available for the current pollution of soil and sediments by these explosives.

In Puerto Rico (PR), significant areas of the Island municipality of Vieques were used by the US Navy in military training exercises for over 60 years. One of the most impacted areas is the Anones Lagoon located at the center of the former Atlantic Fleet Weapons Training Facilities (9). In 2004, a year after the military activities ceased, the US EPA designated the area as a Superfund Site. Only a few studies on the military impacts on Vieques have been conducted by the US EPA and local scientists (10, 11, 12, 13, 14). These studies highlighted the impacts to plants, algae, animals, and public health caused by the presence of metals, explosive compounds and bi-products. Unfortunately, the ecological impacts in the Anones Lagoon by the long-term bombing are still poorly understood.

The study of microorganisms in these atypical sites could reveal unique diversity associated with environmental pollution and could help estimate, better understand and model the biological attenuation processes associated with the area. For instance, certain microbial species can serve as indicators of pollution or an environmental disturbance (15). Microorganisms can also express specific genes to biodegrade or biotransform contaminants into less hazardous compounds. Specific metabolic pathways for degrading toxic compounds generated by military activities have been elucidated, and are often enriched in this type of environment that is hostile to life (16, 17, 18, 19). Genes, including nitroreductases, have been shown to be involved in degrading different explosives (20, 21, 22, 23, 24, 25). Although, these approaches have advanced our understanding of the role of microbes in military impacted sites, all rely on culture-dependent techniques in the laboratory and thus, the relevance of the results obtained from *in-situ* activities and processes remain frequently inaccessible.

Various studies have been conducted using metagenomics analysis to better understand environments such as wastewater treatment plants (26), contaminated freshwater and marine sediments by pesticides (27, 28), and contaminated sediments by oil spills (15). Military sites remaining comparatively much less studied by culture-independent techniques, especially in terms of studying the corresponding microbial communities across time. To our knowledge, there has not been a metagenomics survey of the Vieques military-impacted site or similar tropical lagoon sites. Yet, the relative abundance of microorganisms over time and their gene complement at Vieques could provide new insights into the process of natural attenuation of explosive pollutants or enhanced restoration, and serve as biomarkers for predicting the fate of explosives.

We hypothesized that microbial communities in Anones were enriched in broad adaptive strategies, and perhaps unique biodegradation genes of explosives selected by the long-term history of diverse pollution while increase in microbial diversity could indicate progress of natural attenuation processes. We employed shotgun whole-community metagenomics to test this hypothesis and characterize the genetic diversity and biodegradation genes at the Anones Lagoon. Comparative metagenomics with microbial communities from undisturbed and human-impacted lagoons in Puerto Rico revealed that the Anones sediment communities harbor novel diversity that has apparently contributed to the natural restoration of the site.

## Results

### Site and sample description

The three lagoons used in this study were exposed to different natural or anthropogenic effects. In the case of Guaniquilla Lagoon, The Department of Natural and Environmental Resources of Puerto Rico has since 2002 designated the lagoon as a natural wildlife reserve for native and endemic species (29). On the other hand, Condado lagoon is located in San Juan, which is the capital city (~400,000 inhabitants) and urban area of Puerto Rico. Anthropogenic activities such as urban development with significant sewage discharges has been extensively reported at the site. However, in 2013 the lagoon was designated as an estuarine reservoir (30). Finally, Anones lagoon, the most severely impacted site, was exposed to military training exercises for over 60 years, since it was located in center of the former Atlantic Fleet Weapons Training Facilities (10). At the present time, the lagoon is a Superfund Site designated by US EPA since 2004 and been placed under the jurisdiction of US Fish and Wildlife Service (USFWS). The lagoon is connected to Bahía Salinas del Sur, the bay to the open sea, through a small channel and traces from bullets and bombs are easily spotted at the surface throughout.

Each sediment sample had similar physicochemical characteristics since all lagoons experience similar climatic and edaphic conditions. All samples originated from the surface sediment of each lagoon, as shown in Table 1. In general, pH measurements were nearly neutral, between 7.10 and 7.75, with Anones-2005 being the most alkaline. All samples exceeded the ocean’s typical salinity (~3.5%), except Condado. Heavy metals measurements showed that Condado and Anones-2005 had the highest concentration of lead and cadmium, 15.8μg/g, 0.7μg/g and 34.9μg/g, 0.3μg/g, respectively. Anones-2014 exhibited the highest level for copper, 63.4μg/g, while Guaniquilla had the lowest concentration for all three elements (Table 1). According to the interim freshwater sediment quality guidelines (ISQG) of the Canadian Council of Ministers of the Environments (CCME) (31), Anones-2005 exceeded the cadmium guideline of 0.6 mg/kg, while Anones-2014 exceeded the copper guideline of 35.7 mg/kg. These reference thresholds indicate the possible adverse biological effects on aquatic systems (32)(CCME 1995). Interestingly, Anones-2005 was the only sample in which explosives were detected above the U.S EPA method 8330B limit.

**Table 1.**
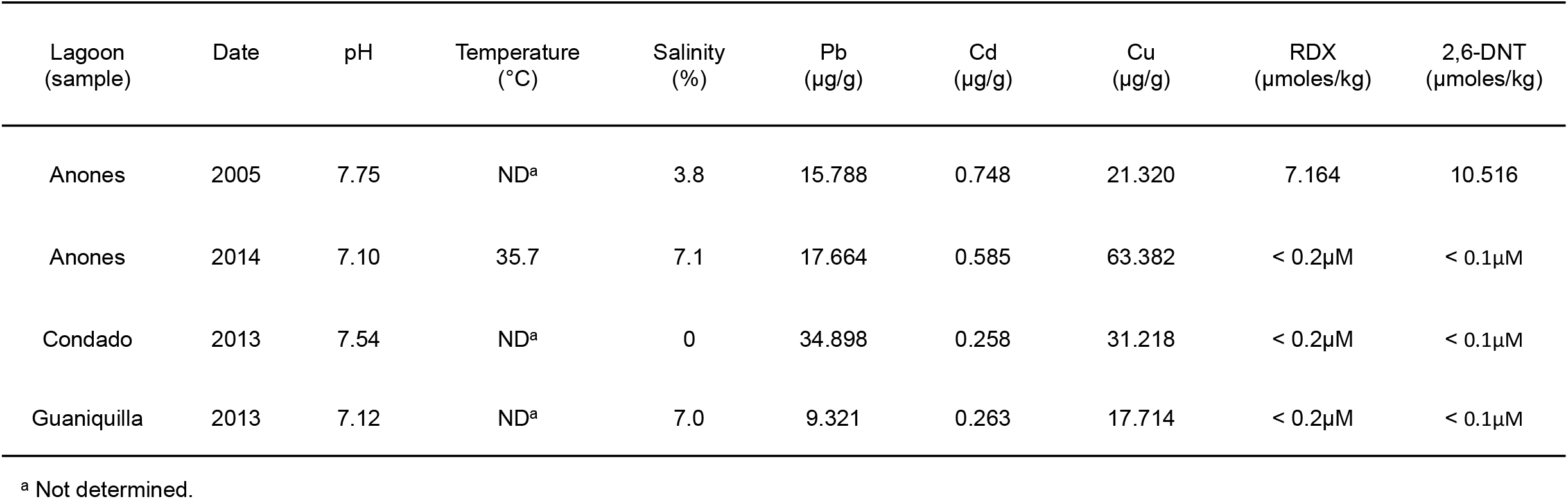
Physicochemical measurements of the sediment samples from each lagoon.

### Microbial community diversity patterns

DNA extraction for each of the four sediment samples (Anones-2005, Anones-2014, Condado and Guaniquilla) was performed in triplicate. The pool of the triplicate samples from each site was sequenced at about 5 Gbp/sample. As typical of metagenomics surveys of complex sediment samples, sequencing did not cover the total DNA diversity sampled (33). Nonpareil 3.0, a database-independent metric of microbial community complexity and alpha-diversity (33, 34), with default parameters showed that the coverage of each microbial community by sequencing was 28%, 33%, 67.5%, and 13% for Guaniquilla, Condado, Anones-2005, and Anones-2014, respectively. Coverage values and Nonpareil curve projections for complete coverage showed that Anones-2005 saturated faster and required the least sequencing effort for complete coverage, while Anones-2014 required the highest (Figure 1A), revealing that Anones-2005 possessed the least diverse microbial community. In particular, the Nonpareil index of sequence diversity (*N_d_*), revealed that Guaniquilla had the highest value *N_d_* = 21.22, closely followed by Anones-2014 at 20.81, whereas Anones-2005 the smallest, *N_d_* =19.68 (note that Nd is a logarithmic scale, thus a difference of 1 corresponds to 10 fold the absolute difference). Comparing the metagenomes from this study to several reference metagenomes determined previously (34), revealed that Guaniquilla and Anones-2014 metagenomes were comparable to other typical sediment samples in terms of coverage and microbial diversity, whereas Anones-2005 was similar to freshwater or heavily impacted environments.

**Figure 1.**
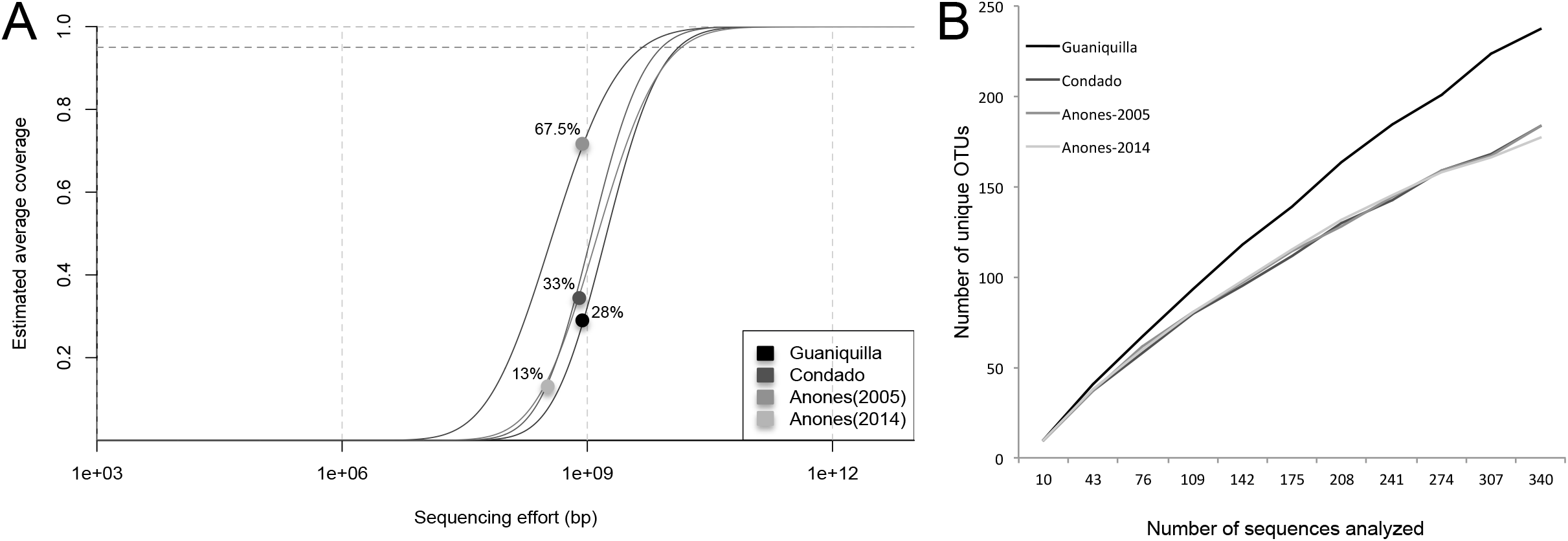
Community (diversity of the three lagoons sampled. (A) Nonpareil 3,0 curves of the Guaniquilla, Condado, Anones-2005 and Anones-2014 lagoon metagenomes, showing the estimated aveoage coverage of the corresponding microbial communities (dots) and the amount of seauencing required to achieve 95% or 99% coverage (dashed lines on the top). Note that curves to the right require more sequencing effort in order to reach high coverage, therefore, the corresponding communities are more diverse. (B) Rarefaction curves of Guaniquilla, Condado, Anones-2005 and Anones-2014 lagoon metagenomes based on 16S rRNA gene fragments recovered. The graph shows the number of unique OTUs per number of sequences analyzed.

### Taxonomic composition and relatedness among the sites

pH and salinity have been known to strongly influence the presence and/or relative abundance of microbial populations (35, 36). In Anones-2005 and 2014, the relative abundance of the two most dominant phyla, *Proteobacteria* and *Bacteroidetes* was similar based on MyTaxa analysis of assembled the metagenomic contig sequences (Figure 2A). In Anones-2005, *Firmicutes* was the third most abundant phylum, making up ~12% of the total; with the remaining phyla collectively making up only 9% of the population. In contrast, Anones-2014 was characterized by high abundance of *Euryarchaeota* (9% of the total), followed by *Firmicutes* (5%), revealing a clear broad taxonomical shift after 9 years, accompanied by an increase in microbial diversity. Further, Anones-2005 was dominated by a few operational taxonomic units (OTUs) based on 16S rRNA gene fragments recovered from the metagenomes (Figure 2B), consistent with a low-diversity microbial community.

**Figure 2.**
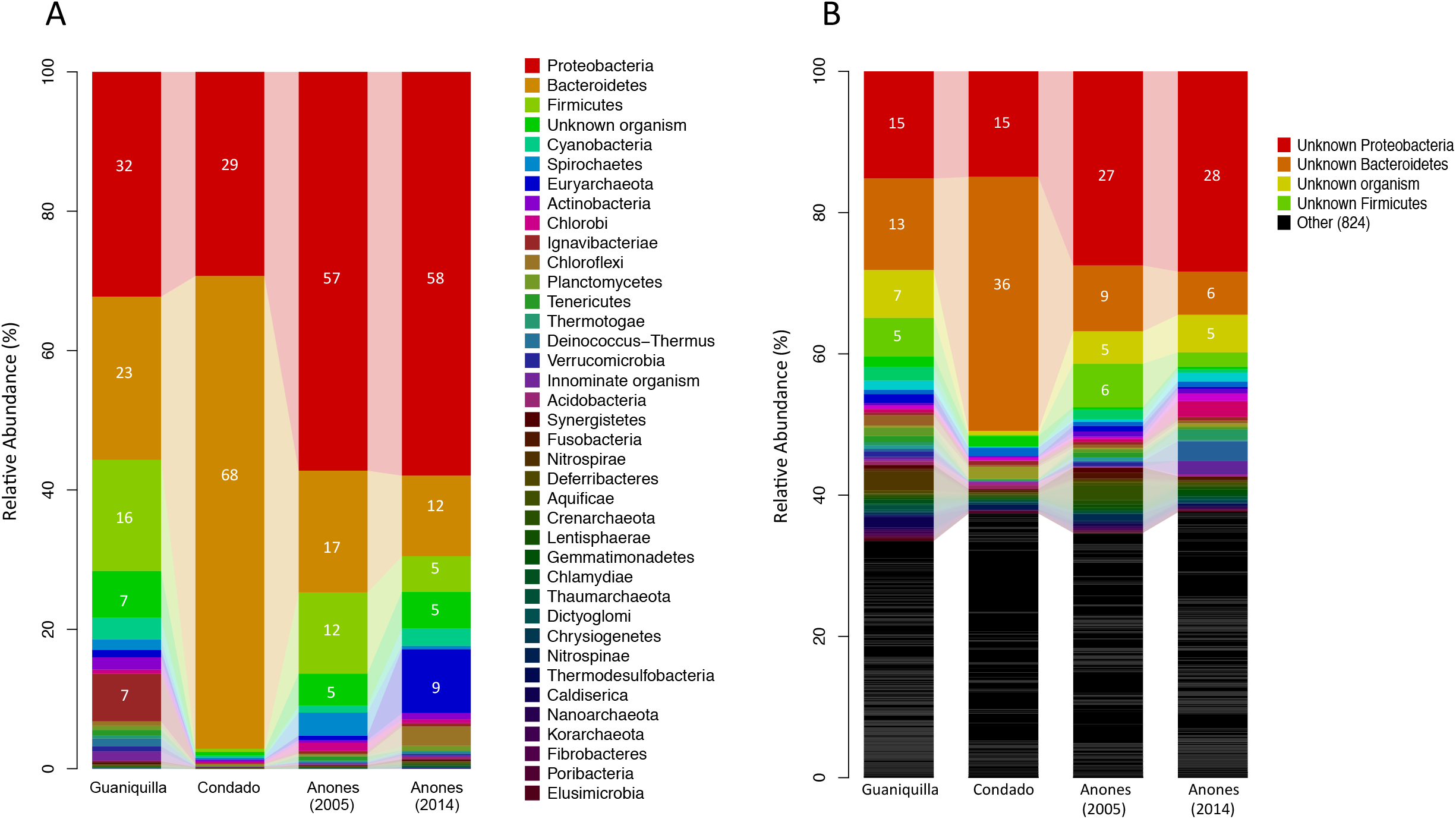
Taxonomic classification of the abundant organisms present in the metagenomes of this study. (A) Phylum-level and (B) genus-level taxonomic classification of the lagoon microbial communities. The graphs are based on MyTaxa analysis of assembled contigs. Numbers indicate relative abundance of each phylum, based on the total reads of the metagenomic dataset (minimum abundance shown is 5%).

The constantly human-impacted lagoon at Condado was dominated by *Bacteroidetes* (68%) and *Proteobacteria* (29%); and only 3% of the metagenomic sequences were assignable to other phyla. Meanwhile, the Guaniquilla lagoon, the least impacted ecosystem by human activity, did not appear to have a dominant phylum with higher than ~30% overall relative abundance. Also, Guaniquilla had the highest abundance of unclassified OTUs (7% of the total) and OTUs assigned to *Ignavibacteriae* (7%), which contrasted with <0.5% *Ignavibacteriae* in the other metagenomes (Figure 2A). Consistent with these findings, the MASH-based distances among the metagenomes showed that Condado was the most distant from either of the Anones samples, reflecting presumably its constant human-induced impact, while Guaniquilla appeared to be more similar to Anones-2014 in complexity (Fig. S2).

To further investigate if the high abundance of the *Bacteroidetes* phylum in Condado lagoon reflected the presence of human-gut associated taxa (due to the location of this lagoon in the city of San Juan), the metagenomic reads were searched against the human gut microbiome IGC reference database (37) for high identity matches (>95% of nucleotide identity). Human gut-associated *Bacteroidetes* genera such as *Bacteroides, Prevotella* and *Porphyromonas* were present in Condado at very low relative abundances (< 0.001%), similar to the other lagoons (Fig. S3). Further, 40% of the detected *Bacteroidetes* belonged to the *Flavobacteria* 3% to *Cytophagia* and 3% to *Shingobacteria* classes, which are typically associated with natural settings such as fresh/saltwater, soil, activated sludge, and compost (38). Hence, it appears that environmentally-adapted populations made up most of the *Bacteroidetes* signal in Condado.

### Biodegradation genes of explosives and heavy metals

In the Anones samples, a significant functional shift was observed between 2005 and 2014 samples (Figure 3). Anones-2014 showed an average decrease of 91.5% in reads mapping to several key genes encoding enzymes involved in the biodegradation of explosives and nitroreductases, including cytochrome P450-like protein (*xplA*), xenobiotic reductase B (*xenB*), xenobiotic reductase A (*xenA*), major oxygen-insensitive nitroreductase (*nfsA*), oxygen-insensitive nitroreductase B (*nfsB*), oxygen-insensitive nitroreductase (*nitA*), oxygen-insensitive nitroreductase (*nitB*), cadmium-transporting ATPase P-type (*cadA*), zinc, cobalt and lead efflux system (*zntA*), Pb(II) resistance ATPase (*pbrA*), copper-exporting P-type ATPase A (*copA*), copper-exporting P-type ATPase B (*copB*) genes compared to Anones-2005; while *xplA, xenB, nfsB, nitA* and *cadA* did not recruit any reads. In fact, Anones-2014 looked more similar to the reference, pristine lagoons in this respect, e.g., only 0.017%, 0.018% and 0.027% of the total reads mapped to the above mentioned genes for Anones-2014, Guaniquilla, and Condado, respectively, contrasting with 0.2% for Anones-2005, i.e., a ~10 fold higher abundance, on average (Table S2). Interestingly, *copA*, a copper-exporting P-type ATPase, appeared to be the gene with the highest number of matching reads in all metagenomes. This finding is consistent with *in-situ* copper concentrations in each sediment lagoon, which were higher than the other two heavy metals assessed.

**Figure 3.**
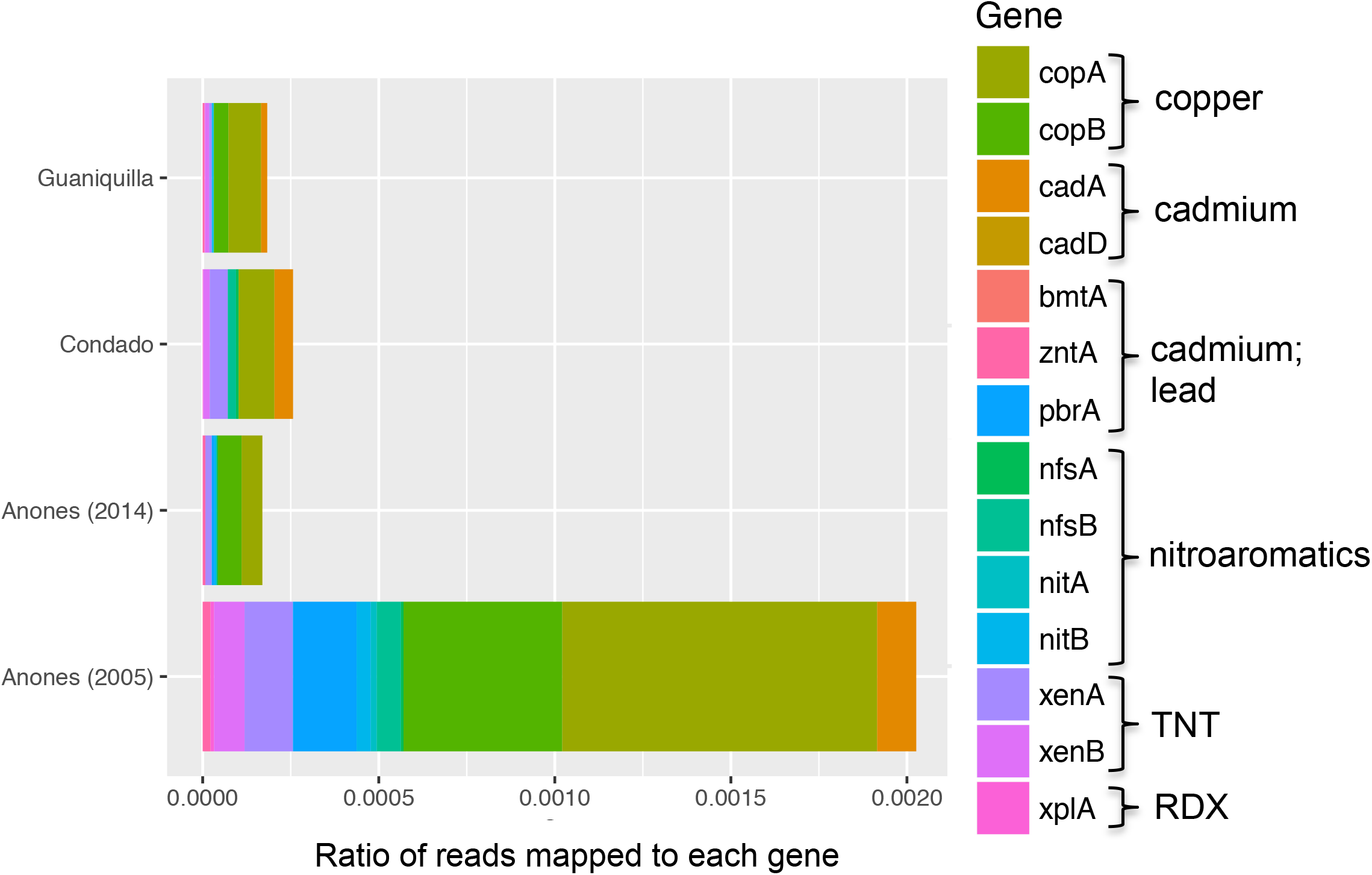
Relative abundance of biodegradation genes involved in explosive and heavy metal (Pb, Cd and Cu) biotransformations in the metagenomes of this study. Reletive abundaeee was estimated as the fraction of metagenomic reads assigned to each specific gene, divided by the number of total reads in each metagenome.

### Novel organisms in Anones 2005

Due to the low complexity of Anones-2005, we were able to recover the genome of five metagenome-assembled genomes (MAGs) representing relatively abundant populations, with high completeness (>83.2%) and low contamination (<10%) using binning techniques (Table 2). The estimated size of these genomes ranger between 2.6 and 4.5 Mbp. Their closest relative in NCBI’s RefSeq prokaryotic genome database showed <50% genome-aggregate amino-acid identity (AAI) and was affiliated with different phyla, indicating that these genome bins (MAGs) represent diverse, novel genera, if not families (39). MAG 2 especially appeared to represent a class-level novel taxon with a low GC (%) content of 31.7% in comparison with the other MAGs (Table S3). Interestingly, all MAGs recovered in Anones-2005 did not appear to be present 9 years later in Anones-2014 (Fig. S4). MAG 3 had a related but distinct microbial population in the Condado metagenome (~90% AAI; Fig. S4). Also, MAG 2 and MAG 7 appear to have a related microbial population in Guaniquilla (Fig. S4).

**Table 2.**
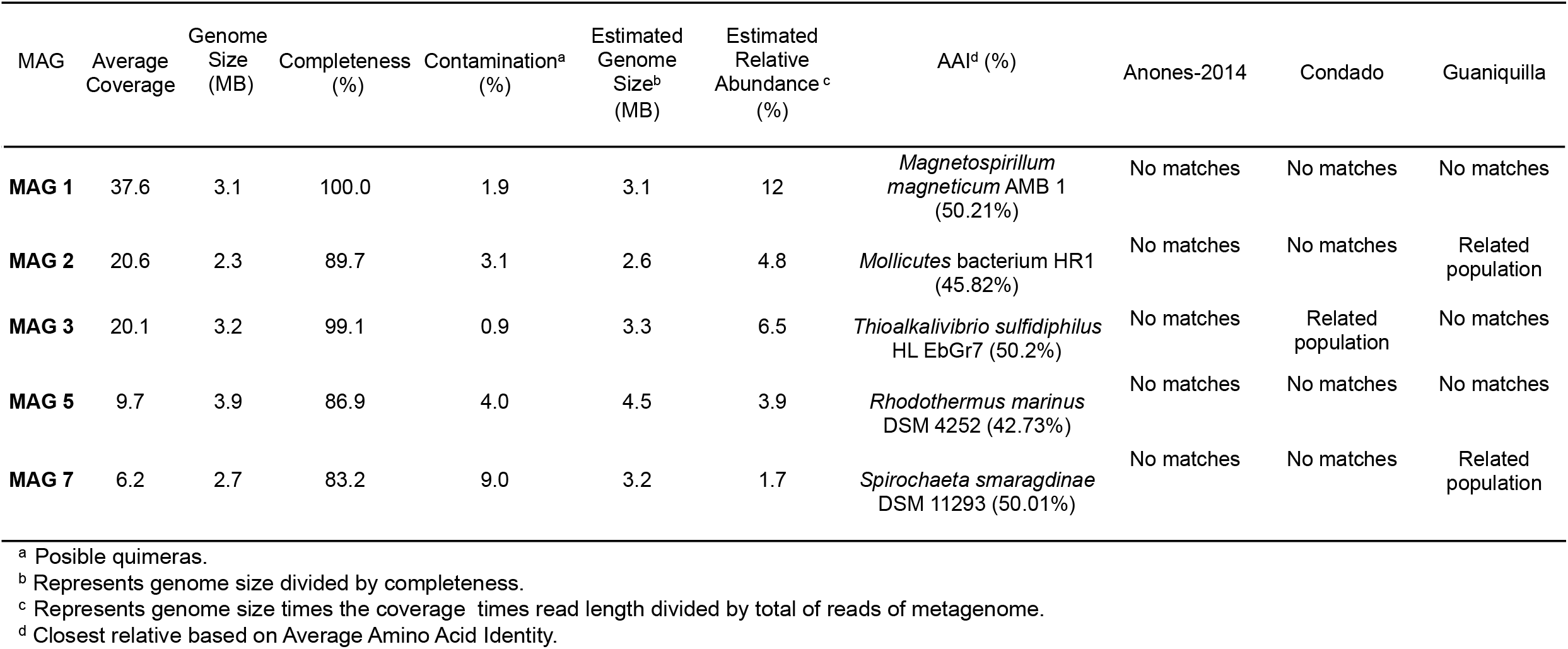
Statistics of the MAGs recovered from the Anones-2005 Lagoon by MaxBin.

MAG 1 (*Rhodospirilaceae* sp., *Alphaproteobacteria*) had a relative abundance of 12% (of the total metagenome) and, together with MAG 3 (*Gammaproteobacteria* sp.) (6.5%), were assignable to the *Proteobacteria* phylum. MAG 2 (*Mollicutes* sp.) (4.8%) was assignable to the *Tenericutes* phylum. *Mollicutes* (MAG 2) and *Bacteroidetes sp*. (MAG 5) represented more deep-branching members of the *Tenericutes* and *Bacteroidetes* phyla, respectively, compared to MAG 1 and MAG 3, based on their best AAI values against RefSeq genomes, and phylogenetic relationships using 57 universal housekeeping genes (Figure 4). All five genomes together accounted for about ~29% of the total metagenome.

**Figure 4.**
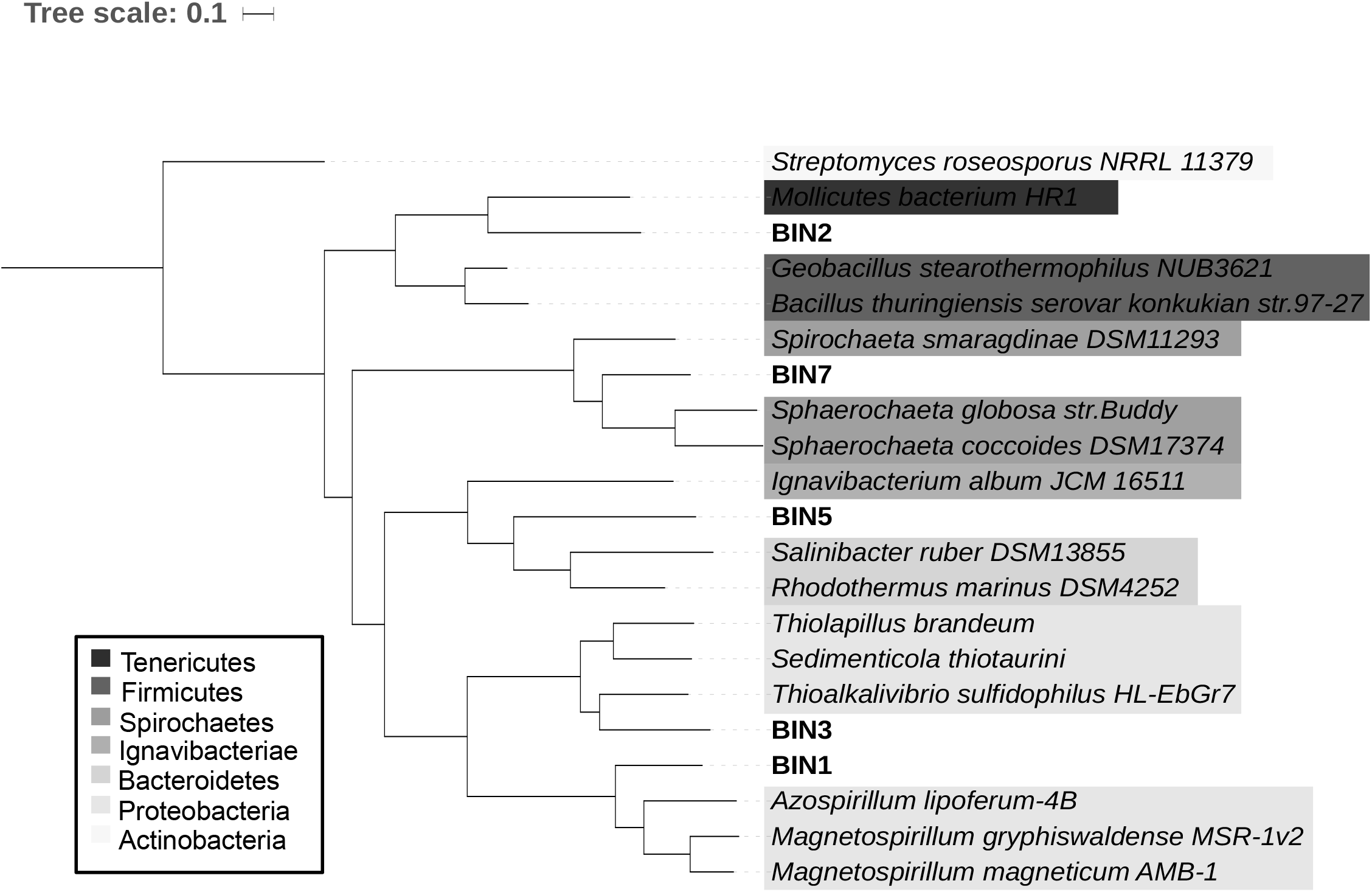
Phylogenetic relationships among; recovered genomes (MAGs) from the metagenomes of this study and selected reference genomes. The phylogeny is based on a maximum likelihood analysis of the concatenated alignment of 57 single copy genes using RAxML and color-coded by phylum.

### Functional description of MAGs

As expected, subsystem categories such as protein metabolism, cofactors, vitamins, prosthetic group, pigments, and amino acid and derivatives were the most abundant pathways in each MAG, followed by DNA and RNA metabolism (Table S4). In addition, each MAG encoded various specialized functions such as: (1) cellular response to DNA damage, (2) response to heat, (3) response to stress, (4) sodium ion transport, (5) SOS response, while no MAG represented photosynthetic bacteria (Figure 5). All MAGs also encoded various manually verified genes with significant homology (e.g., >30% amino acid identity across >70% of the gene length) to genes previously shown to be involved in the transformation and biodegradation of explosives and heavy metals resistance. A brief description of the functional gene content of each MAG follows:

**Figure 5.**
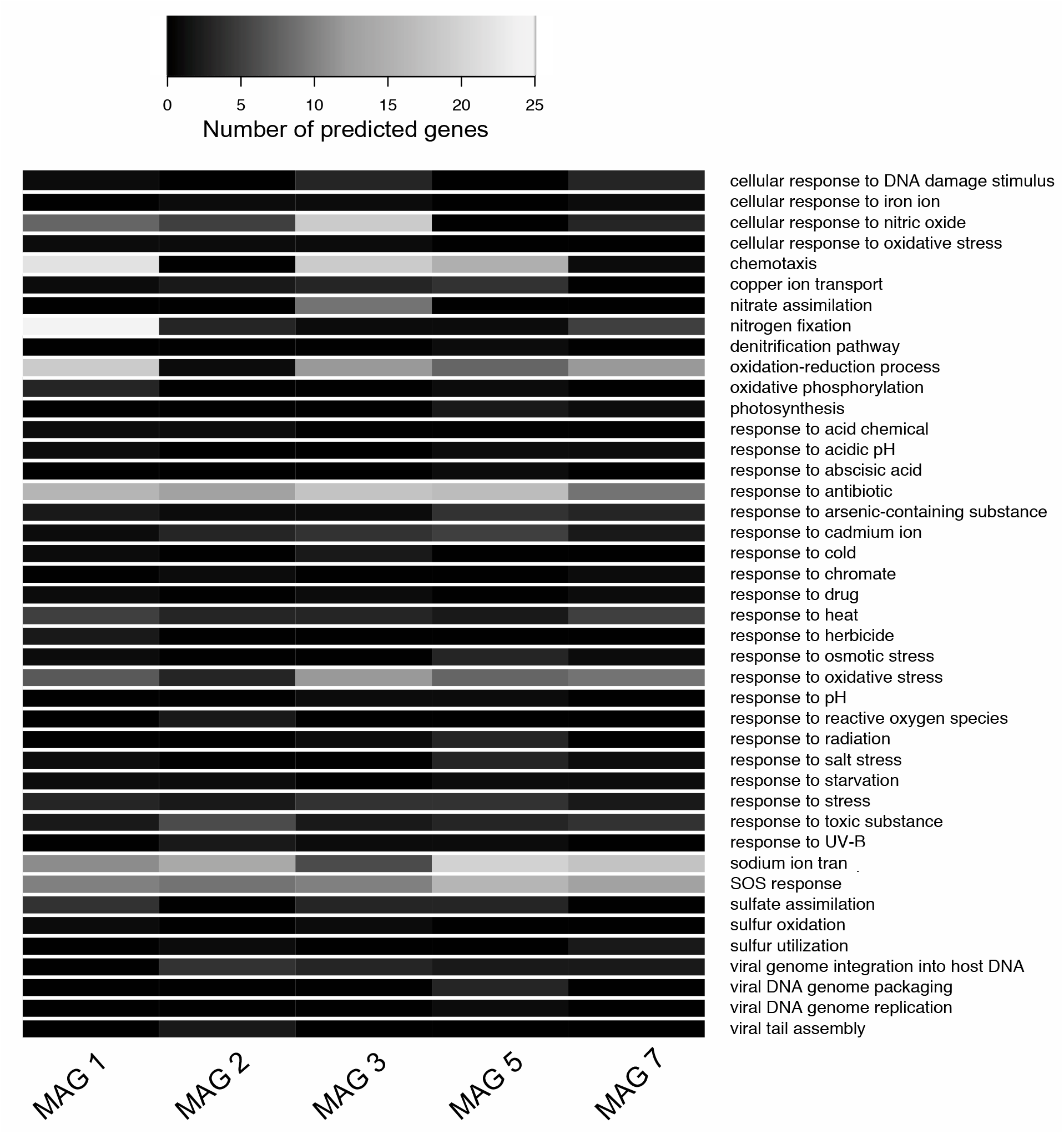
Functional gene annotation of MAGs based on the SwissProt database. Heatmap shows the counts (i. e., number of genes found; see scale bar on top) of gene functional categories (rows) in each metagenome of this study (columns). For each functional category shown, all genes assigned to this function in SwissProt were used to estimate the counts.

### MAG 1 (*Rhodospirilaceae* sp.)

The most abundant MAG appears to belong to the *Proteobacteria* phylum, has a genome size of 3.1Mbp, a GC content of 62.6%, and coding density of ~90%. MAG 1 harbors genes for sulfate assimilation, including adenylyl-sulfate kinase (*cysC*) and phosphoadenosine phosphosulfate reductase (*cysH*) genes. Therefore, this genome is likely from a sulfate-reducing bacterium using sulfate as terminal electron acceptor. Interestingly, MAG 1 is unique among the other four MAGs in harboring the full gene complement (e.g., 25 genes) of a nitrogen fixer, including nitrogenase iron protein 1 (*nifH*). MAG 1 also has genes for chemotaxis, including flagellar motor switch proteins, such as *fliN* and *fliG*, indicating that it is potentially motile. Presence of efflux system proteins involved in response to antibiotics were also detected. In addition, MAG 1 harbors a (predicted) homologous gene to xenobiotic reductase (xenA) and copper exporting (*copA*) (Table S5).

### MAG 2 (*Bacilli sp*.) and MAG 7 (*Spirochaetales sp*.)

MAG 2 was the most deep-branching of all MAGs with respect to the available genomes of isolates in NCBI’s RefSeq prokaryotic database and did not appear to be motile, with a genome size of 2.3Mbp, GC content of 31.7%, and coding density of ~93%. MAG 2 has genes for resistance to antibiotics, reactive oxygen species, and ultraviolet radiation (UV) by homodimerization activity. Viral genome integration proteins, including *int* for integrase functions, appears to be present, indicating vphage predation for this population. Using PHASTER with default parameters (40), only this MAG showed the presence of phage-like proteins, including transposase, portal protein, integrase, head protein and terminase in three different contigs. However, the phage genome was not intact/complete, but most likely represents a remnant prophage. For specific genes of interest, this MAG harbors (predicted) genes associated with cadmium transporter (*cadA*) and copper exporter (*copA*), xenobiotic reductase (*xenA*) and oxygen nitroreductase (*nitB*) (Table S5).

MAG 7 has a genome size of 2.7Mbp, GC content of 50.8%, and coding density of ~93.2%. This genome harbors sulfur utilization proteins such as L-cystine-binding protein (*fliY*) and antibiotic resistance genes (e.g., efflux pump systems). Genes encoding the homologous functions for oxygen nitroreductase (*nitA*), xenobiotic reductase (xenA), copper exporting (*copB*) were also observed (Table S5).

### MAG 3 (*Gammaproteobacteria sp*.)

The second most abundant genome appears to belong to the *Proteobacteria* phylum, has a genome size of 3.2Mbp, GC content of 51.2%, and coding density of ~90%. The presence of nitrate and nitrate reductases indicated that MAG 3 was a denitrifying bacterium encoding primarily facultative anaerobic heterotrophic lifestyle. MAG 3 also harbors genes for chemotaxis (*fliN* and *fliG*) and thus, like MAG 1, appears to be motile. Also present were genes for antibiotic resistance (e.g., efflux pump system), and response to gamma and ultraviolet radiation. For specific genes of interest, MAG 3 appears to carry a gene encoding the XenB protein, which works under anaerobic conditions and less toxic compounds are produced during biodegradation, (50%, that has a high amino acid identity to a previously characterized XenB (41)), oxygen nitroreductase (*nfsB*), and copper exporting (*copA*) genes (Table S5).

### MAG 5 (*Bacteroidetes sp*.)

MAG 5 has a genome size of 3.2Mbp, GC content of 42.2%, and coding density of ~92.5%. It harbors sulfate assimilation genes, including phosphoadenosine phosphosulfate reductase (*cysH);* therefore could be a sulfate-reducing bacterium, likely using sulfate as terminal electron acceptor. It also harbors nitrous-oxide reductase (*nosZ*) gene, i.e., nitrous oxide reductase to atmospheric dinitrogen. MAG 5 also possesses genes for *fliN* and *fliG* genes involved in chemotaxis and thus, likely represents a motile population. Also, present are genes involved in antibiotic resistance (e.g., efflux system), response to radiation, including UV by homodimerization activity, and presence of *int* gene, for viral/prophage integration. Finally, this MAG possesses homologs of oxygen nitroreductase (*nfsA*), copper exporting (*copA, copB*), cadmium transporter (*cadA*), and lead resistance (*pbrA*) genes (Table S5).

## Discussion

The microbial community structure of surface sediment samples from three different coastal lagoons in Puerto Rico was evaluated. Anones was impacted by a major and toxic disturbance; the lagoon was sampled two years after continuous pollution disturbance stopped and then 11 years later. Condado has continuous anthropogenic disturbance due to its proximity to Puerto Rico’s capital city. Finally, Guaniquilla lagoon which is a proxy for a pristine lagoon, since it is a natural reserve. Collectively, our results showed that microbial diversity of the Anones Lagoon during 2005 was negatively affected by military activities since microbial community was ten times or more less complex than the average sediment from reference lagoons or Anones nine years later, and encoded at least 10 times more genes related to biotransformation of xenobiotics and heavy metals. These results indicated that continuously bombing for decades selected for a few populations well-adapted to an environment containing explosive chemicals. Nonetheless, about a decade later, conditions became favorable for bacterial diversity to recover, at least partially. Our findings are consistent with several studies that have documented reduction in microbial community diversity following a major environmental disturbance (42), but also enrichment of specific microbes after exposure to explosives (19, 43, 44, 45, 46, 47).

The substantial microbial community shifts from 2005 to 2014 in Anones also indicated a well-adapted/enriched community to the explosives and under transition in 2005. The major changes observed in microbial composition and diversity nine years after indicates that changes were likely occurring at the end of 2003, when bombing ceased, and the 2005 sample represented an early recovery stages of the microbial community from the use of explosives. Anones-2005 also had the most alkaline pH among the sampling sites, contrary to other military impacted zones with acidic pH (7), which is consistent with a transitional state for the 2005 community. Unfortunately, samples from 2003 or earlier were not available to further corroborate these interpretations. Further, enrichment of *Archaea* has previously been shown to be related to community recovery from an oil spill (15), and *Euryarcheaota* were collectively much more abundant in Anones 2014 vs. 2005. Therefore, it appears that such archaeal populations may represent good indicators of less-polluted ecosystems or recovered communities.

The Anones-2014 microbial community was characterized by a higher number of microbial taxa and lower frequency of biodegradation process genes compared to Anones-2005 or Condado lagoons. The decrease in relative abundance of known genes related to the transformation/biodegradation of explosive compounds between 2005 and 2014 samples strongly indicated that Anones Lagoon has been undergoing natural attenuation. For instance, nitroreductases such as those detected in the Anones-2005 metagenomes and MAGs are known to act by cleaving nitroaromatics rings, including those found in explosives like TNT (22, 23, 24, 25). These findings further confirmed that microbes are capable of faster recovery and adaptation when compared to multicellular organisms (48, 49, 50), and can be used as more sensitive biomarkers of the current state and future projection of ecosystem recovery (51).

Bioinformatics functional prediction of the recovered MAGs from Anones-2005 indicated that the organisms sampled were, at least partially, responsible for the bioremediation enhancement in Anones. MAGs encoded genes related to explosives and heavy metals (e.g., cadmium, copper, and lead resistance genes), including genes for the biodegradation/transformation of nitroaromatics such as TNT and RDX (16, 17, 18), that were absent in other lagoons or Anones-2014. Also, based on AAI values, these organisms represented at least novel families, revealing that disturbed tropical sites by military activities may have selected for novel organisms that need to be studied in more detail both taxonomically as well as functionally for their biodegradation/transformation potential.

The differences in microbial community composition observed in the Anones samples are unlikely to be attributed to seasonal effects or sample heterogeneity. First, our samples represent composite samples of multiple DNA extractions in order to reduce sample-specific patterns and were collected at the same time of the year (summer). Further, previous studies have shown that bacterial communities inhabiting sediments of tropical coastal lagoons do not show strong seasonal patterns (52, 53), consistent with relative small seasonal variations in temperature in the tropics.

In conclusion, this study has revealed the potential functions and organisms associated with transformation/biodegradation of explosives and resistance of heavy metals in a former bombing range. Even though our results indicated that natural attenuation has been occurring since 2005 in Anones, it is important to note that it only involved the surface of the lagoon’s sediment. Thus, the structure of microbial communities residing at deeper sediments remains uncharacterized. Several questions also remain to be addressed in future studies; most notably, how much the identified microbial populations contributed to natural attenuation and if their activities are adequate for complete bioremediation of a site constantly disturbed for over 60 years by military activities. Tracking temporal shifts after the disturbance, coupled to *in-situ* rate measurements, could be a useful approach to better quantify the role of benthic microbes for natural attenuation of explosives and other environmental pollutants, and provide biomarkers for better modeling the attenuation process and predicting the toxic effects of specific chemical compounds. The genes and genome sequences recovered here can also provide reference points for future experiments related to the remediation of Anones or other contaminated sites, e.g., by providing sequences for qPCR assays.

## Materials and Methods

### Sampling

Soil samples from the sediment surface (0-5cm), where heavy metals and explosives residues were mainly deposited (6, 7), were collected in Corning^®^ 50mL centrifuge tubes from Anones at two time points (October 2005 and June 2014). Similarly, sediment surface soil smaples from the pristine Guaniquilla and urban impacted and Condado lagoons (Fig. S1) were collected in June 2013. All samples were stored at 4°C until further analyzed. Temperature measurements were taken *in-situ* by immersion of a mercury thermometer in sediment; a salinity refractometer was used to measure salinity of water above the sampled sediment, and pH measurements were taken on the samples in the laboratory with a pH meter (ATI Orion Model 230A).

### *Heavy Metal* and *Explosive Concentrations*

To measure heavy metals an acid extraction was done, approximately 3g of homogenous sediment sample was incinerated for 3hrs at 600°C in ceramic crisols. After incineration, samples were pulverized with approximately 1mL of concentrated HCl and transferred to a 250mL beaker by rinsing with 1mL of HCl for a total of three washes in order to transfer all the sample to the beaker. One mL of nitric acid was added to each sample while warming samples without drying, for a total of two washes and a last wash of 3mL concentrated HCl. By the end of the washes, the sediment was white in color. The acid extraction was then filtered through a Whatmann filter #40 and 5% HCl added for a total of 100mL of sample. Lead (Pb), cadmium (Cd), and copper (Cu) were measured by atomic absorption spectrophotometer (Perkin Elmer Model AA100). Concentrations for RDX and 2,6-DNT were measured by High Performance Liquid Chromatography (HPLC) according to U.S EPA method 8330B (54).

### DNA extraction protocol

Metagenomic DNA extraction was performed with the modified DNA extraction protocol Method 2 using the QIAamp DNA Micro Kit from QIAGEN^®^ (55). Briefly, approximately 0.5g of homogeneous sediment sample was subjected to a two-step cell lysis. First, a chemical lysis was performed by addition of an enzyme cocktail as follows: Mutanolysin (1,500 u/ml), Lysostaphin (510 u/ml), and Lysozyme (10mg for 2ml), plus a lysis buffer [0.5M EDTA and of 1M Tris (pH 8.3)] and incubation at 37°C for 1h with rotation to mix. Second, a physical lysis was performed with approximately 0.22g of each bead (0.1mm glass and 0.5mm ziconia/silica beads) and 600μl Phenol:Chloroform at 3,500rpm for 1:30 min. After centrifugation at 3,000rpm for 5min and transferring of the supernatant to a new microtube, AL Buffer (QIAGEN kit) and 100% EtOH were added to the supernatant. The rest of the protocol followed the QIAamp DNA Micro Kit manual from step 17 to protocol end.

### High-throughput sequencing and sequences processing

Community DNA (libraries) was sequenced using an Illumina MiSeq reagent V2 kit for 500 cycles (2 x 250 bp paired end run) on an Illumina MiSeq instrument (located in the School of Biological Sciences, Georgia Institute of Technology). Prior to sequencing, DNA sequencing libraries were prepared using the Illumina Nextera XT DNA library prep kit according to manufacturer’s instructions except the protocol was terminated after isolation of cleaned double stranded libraries. Library concentrations were determined by fluorescent quantification using a Qubit HS DNA kit and Qubit 2.0 fluorometer (ThermoFisher Scientific) and samples run on a High Sensitivity DNA chip using the Bioanalyzer 2100 instrument (Agilent) to determine library insert sizes. Adapter trimming and de-multiplexing of sequenced samples was carried out on the MiSeq instrument. Raw metagenomic reads were trimmed using Solexa Q4 (56). Each resulting trimmed pair-end read was merged together with its sister read, when overlapping, using PEAR with default parameters (57) (Table S1).

### Metagenome coverage and de-novo assembly

Nonpareil, which assesses coverage of extracted community DNA by sequencing based on the frequency of unmatched reads in the metagenomic datasets (35, 58), was used determine the level of microbial community coverage and sequence diversity. In the present study, Nonpareil 3.0 was used, which represents a faster, k-mer-based method than the original Nonpareil, and in addition, provides an estimation of the alpha-diversity of the sample (58). Assembly of the metagenomic reads was performed with IDBA 1.1.1 (59) with a minimum k-mer value of 35 and a maximum value of 75. The k-mer size that resulted in the highest number of assembled reads was selected for each metagenome.

### Taxonomic classification of DNA sequences and estimation of in-situ relative abundance

Taxonomic classification was assessed in two ways: (1) assembled contigs or binned genomes were classified using the stand-alone MyTaxa analysis (60) and reported at the Phylum level; (2) 16S rRNA gene-encoding reads were identified using Parallel-META 2.0 (61) followed by QIIME 1.9.0 (62) for taxonomic classification and the top 75 most abundant genera reported.

Trimmed reads were mapped on predicted genes of contigs using BLAT in order to assess relative abundance of the gene or population bin, based on a minimum cut-off for a match of 97% nucleotide identity. Finally, a distance matrix was developed to estimate sequence relatedness between metagenomes based on MASH distance analysis (63) at the whole metagenome level, and visualized using the PCA plot function of the QIIME principal_coordinates.py script (62).

### Gene prediction and functional annotation of biodegradation genes

Gene prediction was performed with MetaGeneMark (64) using trimmed metagenomic reads or assembled contig sequences as input. Predicted genes were compared against a manually curated *in-house* database of biodegradation genes using BLAST (65) for complete alignment, conservation of functional domains and at least 30% amino acid identity (minimum bitscore cutoff of 60 for a match). The database included genes related to explosives (*xplA, xenB, xenA, nfsA, nfsB, nitA, nitB*) and heavy metals resistance for Pb, Cd and Cu (*bmtA, cadA, zntA, pbrA, cadD, copA, copB*).

### Recovering population genomes by binning

Assembled contigs for each metagenome were binned into MAGs using MaxBin software (66) in order to obtain whole or partial genomes. For each MAG, coverage, genome size, completeness, and contamination were estimated using the HMM.essential.rb script as implemented in the Microbial Genomes Atlas (MiGA) webserver (67). Average Amino Acid identity (AAI) values (68) against the RefSeq prokaryotic genome database from NCBI (https://www.ncbi.nlm.nih.gov/refseq) were also computed by the MiGA webserver.

To further improve the quality of the recovered MAG sequences, the likely phylogenetic origin of the contigs was evaluated as follows: every gene encoded by the contigs of a MAG was searched against the RefSeq prokaryotic genome database for its best match. Contigs that provided matches to different taxonomic families or provided highly divergent AAI values were manually removed from the MAG. Phylogenetic relationships of the resulting MAGs and their best-three matching RefSeq genomes were determined based on their AAI values as well as sequence alignment of 57 essential genes shared by all genomes using RAxML (69) and were checked for consistency. Functional analysis for each MAG was performed with SEED and Swiss-Prot database for more specific functional prediction. Predicted genes were also compared to manually curated biodegradation genes related to explosives and heavy metals resistance as described above.

The raw sequences of each metagenome are available in the Sequence Read Archive (https://www.ncbi.nlm.nih.gov/sra/SRP156313) under Bioproject “Vieques metagenomes” (PRJNA483958) and accession numbers: SAMN09754619, SAMN09754620, SAMN09754621, SAMN09754622. MAG sequences are available through http://enve-omics.ce.gatech.edu/data/vieques

## Acknowledgments

Special thanks to Cacimar Zenón for his help during the sampling trips to Vieques, Elba Díaz for advice on chemical analysis, and Casa Pueblo in Adjuntas for support during the project. Thanks also to Juliana Soto, Luis Miguel Rodríguez-R and Carlos Rodríguez Minguela for helpful discussions related to the analysis of the metagenomic datasets. This research was supported by U.S. National Science Foundation (award 1241046).

